# Experimental evolution of independent genetic pathways for resistance to *Pseudomonas aeruginosa* pathogenicity within the nematode *Caenorhabditis remanei*

**DOI:** 10.1101/484998

**Authors:** Heather Archer, Patrick C. Phillips

## Abstract

Pathogenic host-microbe interactions can result from continuous evolution of a host’s ability to resist infection and a pathogen’s ability to survive and replicate. *Pseudomonas aeruginosa* is a versatile and opportunistic pathogen, ubiquitous in the environment, and capable of damaging plants, vertebrates, and invertebrates. Previous studies in nematodes suggest that the pathogenic effects of *P. aeruginosa* can result from multiple distinct pathways: a toxin-based effect that kills within a few hours and a generalized virulence that kills over the course of multiple days. Using experimental evolution in the highly polymorphic nematode *Caenorhabditis remanei*, we show that nematode resistance to the two modes of pathogenesis in *P. aeruginosa* evolves through genetically independent pathways. These results demonstrate that multiple virulence patterns in a pathogen can result in multiple responses in the host, and the genetic lines established here create resources for further exploration of the genetic basis for resistance to *P. aeruginosa*.

## INTRODUCTION

Pathogenic host-microbe interactions can be likened to an ongoing battle, a continuous evolution between the host’s ability to restrict infecting microbes and the pathogen’s ability to survive and replicate. If an important long-term goal of health research is to sway the battle in favor of hosts, then it is fundamentally important to understand the genetic mechanisms underlying this exchange. As a model system, *C. elegans* is uniquely tractable for the whole-organism study of pathogen interactions. In addition to being a bacteriovore, it is susceptible to many of the same pathogens that sicken and kill mammals and humans (O’Callaghan & Vergunst, 2010; Irazoqui, et al., 2010; Feinbaum, et al., 2012; Arvanitis, Glavis-Bloom, & Mylonakis, 2013). Additionally, these worms have interacted with and adapted to bacteria for greater than 600 million years (Irazoqui, et al., 2010). However, the hermaphroditic mating system of *C. elegans* has resulted in greatly reduced genetic and phenotypic diversity (Andersen, et al., 2012; Phillips, 2012; Teotónio, 2017). This is of particular note because most previous studies examining the genetic basis of pathogen response in nematodes have been conducted using *C. elegans*, with a common conclusion being that the amount of genetic diversity limits evolutionary diversification and the resultant functional bioactivity (Schulenburg & Ewbank, 2004; Barriére & Félix, 2005; Schulenburg & Boehnisch, 2008). Further, it has been shown that coevolution of a host organism with pathogenic bacteria drives self-fertilizing populations to extinction while selecting for increased outcrossing in mixed mating populations (Morran, et al., 2011; Slowinski, et al., 2016). Presumably this arises out of a selection-imposed necessity for allelic and phenotypic variation. (Schulenburg & Ewbank, 2004).

Phenotypic variation induced via mutagenic agents has provided many valuable models of host-pathogen interactions and disease mechanics (*C. elegans* deletion mutant consortium, 2012; Boulin & Hobert, 2013). This blunt force approach, however, frequently identifies only the most severe mutations, dominated by those which completely abrogate or grossly impair gene function (Albertson, et al., 2009). This is in stark contrast to the typical trajectory of novel mutations in natural populations where natural selection and genetic drift lead to an accumulation of mutations with small to moderate effects (Rockman, 2012; Noble, et al., 2017; Teotónio, et al., 2017). Models utilizing natural variation in genetic pathways can reveal physiological mechanisms that modify pathogen interactions and disease which would likely be missed using a mutagenesis approach and evolution itself can be used to select for alleles conferring a desired phenotype (Kammenga, et al., 2008; Teotónio et al., 2017; Gao, et al., 2018; Hahnel, et al., 2018).

To capitalize on natural variation, we have created a new model system using *Caenorhabditis remanei* as host and *Pseudomonas aeruginosa* as pathogen. *C. remanei* is a nematode bacteriovore akin to *C. elegans*. Like *C. elegans* it is easily maintained and manipulated in lab environments, is transparent so that phenotypes can be easily scored, survives indefinitely when frozen, and has a short generation time that readily facilitates multi-generational examination of evolutionary genetic responses to controlled environments. Additionally, it possesses tractable genetic architecture (Fierst, et al., 2015). In contrast to *C. elegans*, however, *C. remanei* is an obligately outcrossing species. The mode and tempo of its population demography is distinct from that of hermaphroditic nematode species, and the resulting amount of genetic diversity present in natural populations is extraordinarily high relative to *C. elegans* (Graustein, et al., 2002; Jovelin, Ajie, & Phillips, 2003; Cutter, Baird, & Charlesworth, 2006).

*Pseudomonas aeruginosa* is a highly virulent and opportunistic mammalian pathogen that is also damaging to plants and invertebrates such as *C. remanei*. It is a frequent cause of death for individuals with cystic fibrosis, severe burns, and other forms of compromised immunity (Tan, Mahajan-Miklos, & Ausubel, 1999). *Pseudomonas aeruginosa* is ubiquitous in soil but possesses a highly variable metabolism enabling it to survive in a wide variety of environments and to produce environment-specific virulence factors (Mahajan-Miklos, et al., 1999; Tan, et al., 1999; Kirienko, et al., 2014). Presumably, the shared ecology between *P. aeruginosa* and *C. remanei* has led to historical host-microbe encounters between these organisms resulting in the presence of natural genetic variation at a relevant scale.

In the lab environment, *P. aeruginosa* can kill *C. elegans* in either a slow infectious process or by a toxin-mediated mechanism depending on the strain and culture conditions (Tan, et al., 1999; Mahajan-Miklos, et al., 1999). Infectious conditions require *P. aeruginosa* to be grown on relatively low osmolarity nematode growth medium (NGM) and have a slower rate of death, whereas the toxin mode of pathogenesis requires a high osmolarity medium and has faster death kinetics. Observations indicate that on infection inducing media bacteria begin accumulating in the intestine of *C. elegans* approximately 36 hours after feeding begins (Tan, et al., 1999). Worms removed from infectious *P. aeruginosa* within 12 hours of exposure or placed on heat-killed bacteria with low osmolarity media do not appear to exhibit any reduction in fitness (Tan, et al., 1999). Under high osmolarity conditions, *P. aeruginosa* produces diffusible toxins and killing does not require contact with live bacteria (Mahajan-Miklos, et al., 1999). More recently, toxin-based killing has been shown to be mediated by production of phenazine-1-carboxylic-acid (PCA) in a pH dependent manner (Cezairliyan, et al., 2013). PCA is produced by all pseudomonads capable of making phenazines and promotes the formation of biofilms by increasing the amount of available ferrous [Fe(II)] iron (Wang, et al., 2011).

The existence of different modes of pathogenesis with different kinetics, each requiring a unique environment, suggests that *P. aeruginosa* may have evolved multiple strategies for host infection as a response to host defenses. This may indicate that host defense mechanisms respond to pathogen damage signatures rather than recognition of the microbe itself. If a microbe-associated molecular pattern underlies the host evolutionary response, then populations historically interacting with a pathogen in a specific environment would also display resistance across novel environments. However, if damage-associated patterns activate host defenses then host-pathogen interactions in novel environments will lack cross-resistance and be independent when modes of pathogen damage are environment specific. To assess the evolutionary response of a natural population of *C. remanei* to *P. aeruginosa*, a 40-generation adaptation study under infectious and toxin-based pathogenesis conditions, as well as their combination, was performed. The results from this work show that at least some of the components of these response pathways are independent from one another, suggesting recognition of damage patterns participates in initiating the immune response in *C. remanei*, and that experimental evolution can be a powerful tool for exploring the genetic basis of pathogen resistance within natural populations.

## RESULTS

### *P. aeruginosa* is pathogenic to *C. remanei*

When exposed to *P. aeruginosa* PA14 under infectious conditions, *C. remanei* recently collected from nature showed no obvious symptoms or increased mortality after 24 hours of exposure. Continued exposure results in a gradual cessation of pharyngeal pumping and loss of mobility followed by a significant increase in mortality by day 3 (Figure 1). Differences between PA14 and the standard bacterial strain used as a reference control (*E. coli* OP50) are most pronounced after 5 days, with only half of the population exposed to *P. aeruginosa* remaining at that time (Figure 1). For simplicity, further tests of infection-induced mortality focus on this 5-day time point.

**Figure 1.**
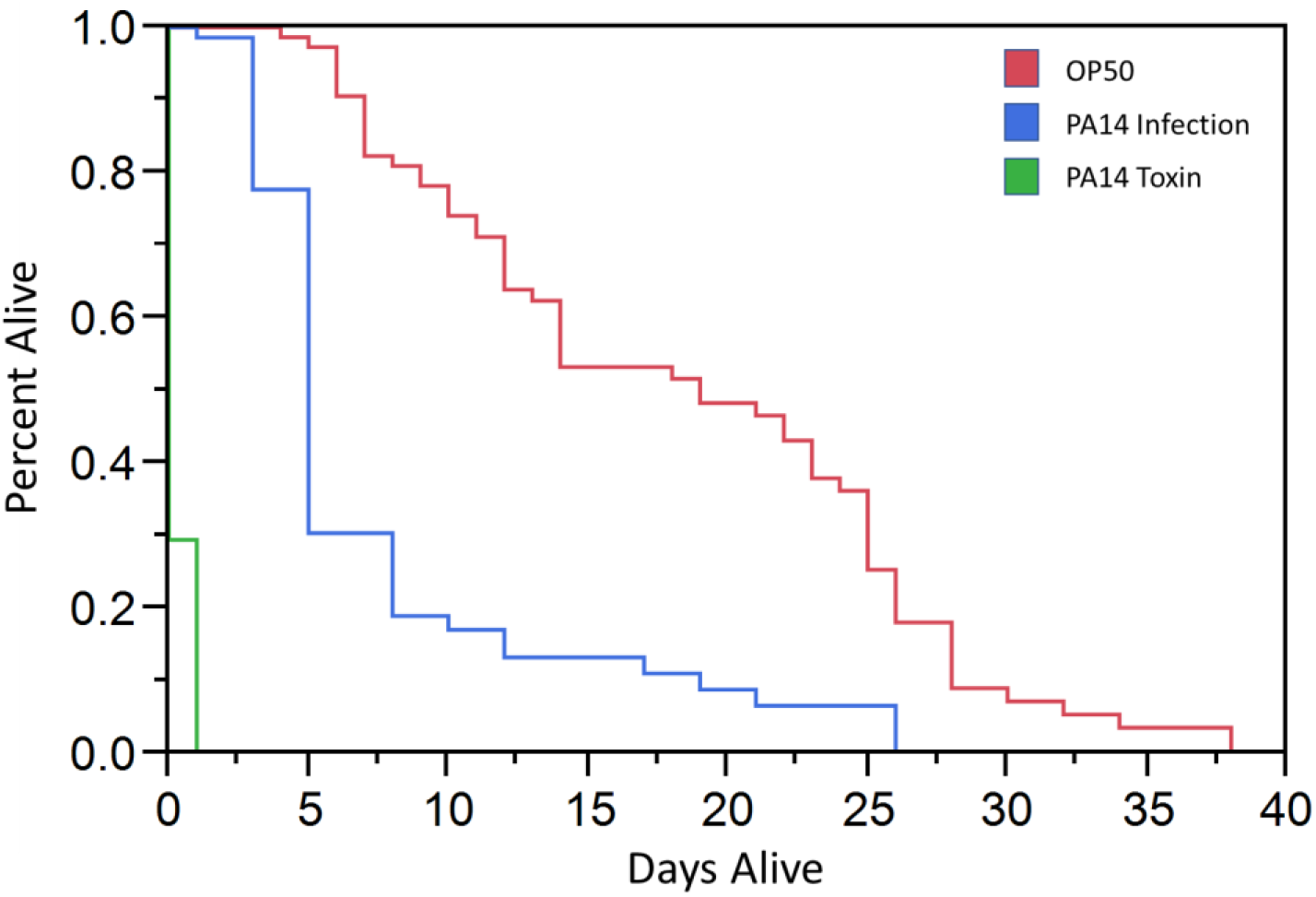
Survival kinetics of ancestor population (PX443) in control (*E. coli* OP50 grown on NGM) and experimental conditions (*P. aeruginosa* PA14, blue = low osmolarity infectious environment, green = high osmolarity toxin producing environment).

*C. remanei* populations are unable to persist for any significant length of time when continuously exposed to *P. aeruginosa* that have been induced to produce the diffusible toxin, showing obvious signs of stress within 4 hours, with approximately 70% of the population dying within 1 day of exposure (Figure 1). Based on these responses, subsequent assessment of toxin-killing resistance consisted of an initial four-hour exposure to the toxin-producing environment, followed by an assessment of individual viability 24 hours after transfer to control conditions.

### Resistance to *P. aeruginosa* is an evolvable trait

After 40 generations of experimental evolution in either infectious or diffusible toxin environments, the resulting *C. remanei* populations respectively showed 16% and 48% increases in resistance to *P. aeruginosa* relative to the ancestral population (Figure 2A; infection, 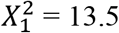, p = 0.0002; toxin, 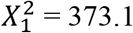, p < 0.0001). As there were no significant differences among replicates (infection, 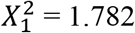, p = 0.1820; toxin, 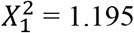, p = 0.2743), data was pooled across replicates for all subsequent analyses (see Supplemental Table 1 for replicate specific information).

**Figure 2.**
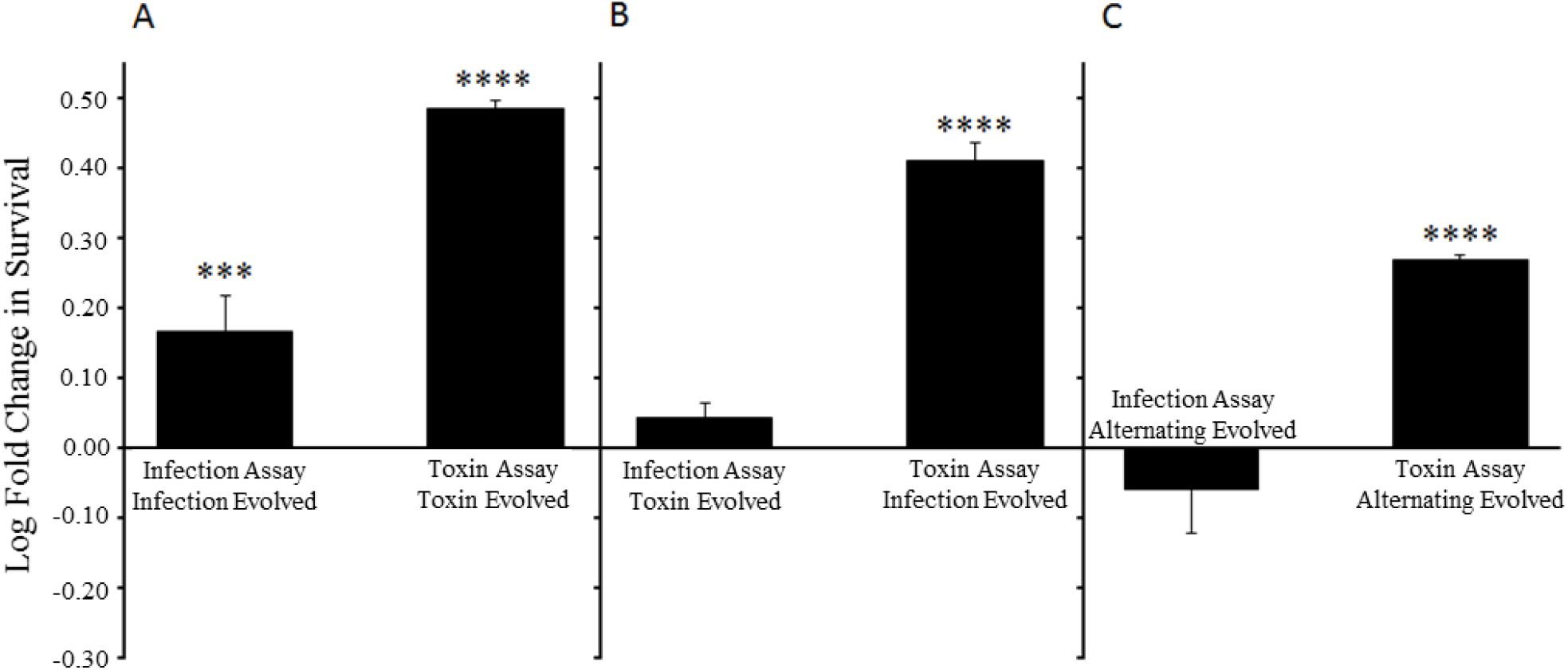
Log fold change in survival relative to ancestral population of *C. remanei* after 40 generations of adaptation. A) Respective environments - infection evolved, 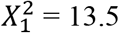, p = 0.0002; toxin evolved, 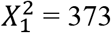, p < 0.0001. B) Reciprocal environments – infection evolved, 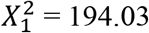, p < 0.0001; toxin evolved, 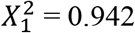, p = 0.3317. C) Alternating population – infection assay, 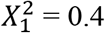, p = 0.5464; toxin assay, 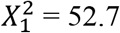, p < 0.0001.

### Resistance evolved in one environment does not necessarily confer resistance to another

It is possible that adaptation within a fixed pathogenesis regime could lead to a correlated increase in resistance in the other regime via shared responses, a negative correlated response due to tradeoffs in modes of resistance to each environment, or independent evolution to each mode with no correlated effects across environments. To distinguish among these possibilities, we tested resistance of populations evolved under one selection regime to the pathogenic effect found in the other regime. We find that individuals evolutionarily adapted to the toxin-producing environment do not display any increase in survival when placed on infection media (Figure 2B, 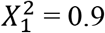, p = 0.3317). In stark contrast, survival in toxin-producing conditions increased significantly for the populations adapted to infection (Figure 2B, 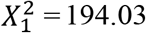, p < 0.0001), although this increase is still somewhat lower than that observed for the toxin-evolved population (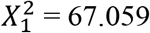, p < 0.0001). Thus, we find asymmetry in the correlated responses with both negative and positive correlations being displayed, depending on the selection environment.

### Evolution under alternating infectious and toxin-producing environments

To further test whether adaptation to one mode of pathogenesis generates a tradeoff in the other, we specifically moved populations between the infection and toxin regimes over the course of 40 generations of adaptation. Similar to the lines evolved with exposure to the toxin environment only, lines evolved under an alternating exposure to infection and toxin display no significant increase in resistance to the infectious environment (Figure 2C, 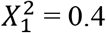, p = 0.5464). However, in the toxin-producing environment, these alternatingly evolved populations have a significant increase in resistance compared to the ancestor (Figure 2C, 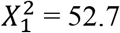, p < 0.0001), although the degree of resistance is lower than all other experimentally evolved populations (Supplemental Figure 1).

### Resistance is also a byproduct of adaptation to the laboratory

Although infection evolved populations survive toxin-mediated killing significantly better than ancestral individuals (Figure 2B), their resistance to the toxin-producing environment is not substantially different from lab adapted controls simply raised on *E. coli* (OP50) (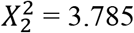, p = 0.1507; Supplemental Figure 2). This indicates that a portion of the mechanism underlying resistance to the toxin phenazine-1-carboxylic acid arises from exposure to standard lab conditions and is independent of interaction with both *P. aeruginosa* and the osmolarity of the media. Importantly, controls evolved under high-osmolarity conditions but with the standard lab food OP50 in place of PA14, do not significantly differ in toxin or infection resistance from controls evolved under standard low-osmolarity lab conditions (Figure 3, 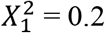, p = 0.6854, and 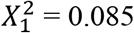, p=0.77, respectively). Although adaptation to standard laboratory conditions appears to benefit resistance to the toxin, it appears to have a mildly negative influence on resistance to the infectious environment (Figure 3, 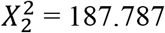, p < 0.0001, and 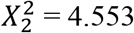, p = 0.10, respectively).

**Figure 3.**
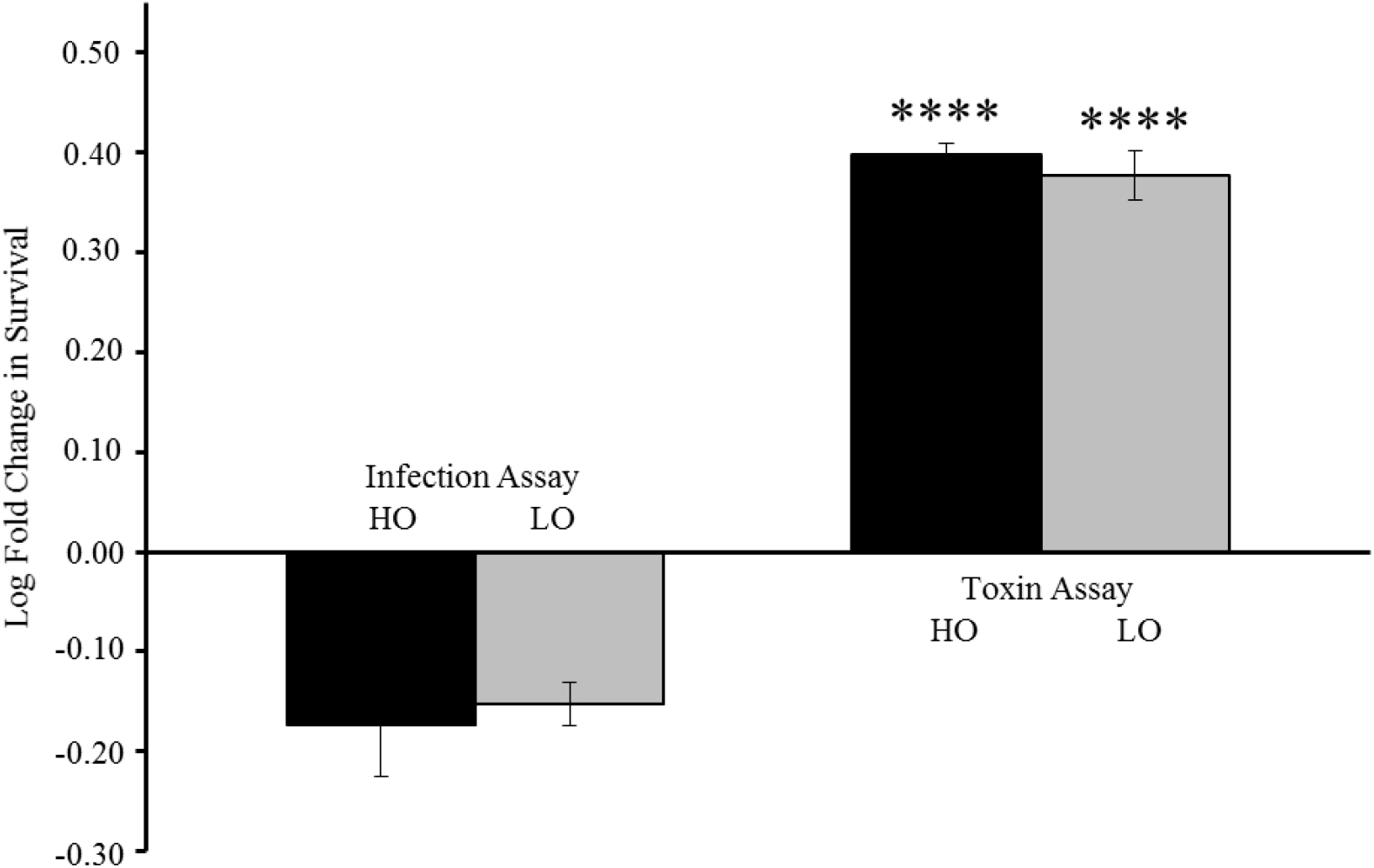
Log fold survival relative to ancestor of lab-adapted (OP50) control lines (high osmolarity control (HO), infection assay, 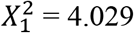, p = 0.05, toxin assay, 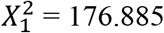, p < 0.0001; low osmolarity control (LO), infection assay, 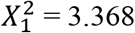, p = 0.07, toxin assay, 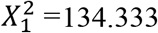, p < 0.0001).

## DISCUSSION

### Contrasting kinetics of pathogenicity in the ancestral *C. remanei* population

In infectious and toxin-mediated killing assays conducted using *C. elegans*, consumption of live bacteria is required for eventual death caused by infection via proliferation of alive cells. In contrast, *P. aeruginosa* grown on PGS media produce phenazine-1-carboxylic acid, a diffusible toxin that leads to mortality in a much shorter timeframe (Tan, et al., 1999; Cezairliyan, et al., 2013). The kinetics of pathogenicity in *C. remanei* appears to be highly like the kinetics seen previously in *C. elegans*, where mortality can also be generated via two distinct mechanisms (Figure 1). In the infectious environment, exposure of *C. remanei* to *P. aeruginosa* for a minimum of 24 hours is required for pathogenesis to manifest itself, with significant mortality at 72 hours of continuous exposure. Under the toxin-producing conditions, only four hours of exposure achieves 70% mortality at the 24-hour time point (Figure 1).

Studies of *P. aeruginosa* mutants with attenuated killing abilities in *C. elegans* has shown that many of the bacterial genes underlying pathogenicity on the low- or high-osmolarity media are specific to that environment, primarily producing phenazines in a relatively narrow and largely non-overlapping range of conditions (Mahajan-Miklos, et al., 1999; Tan, et al., 1999; Cezairliyan, et al., 2013). Pathogenicity in the infectious and toxin-mediated environments clearly arises from distinctly different genetic and biochemical mechanisms. This suggests the possibility there are different modes of pathogenicity from the pathogen’s perspective and distinct genetic mechanisms underlying the response to these forms of pathogenicity from the host’s perspective. However, it is also possible that the basis of resistance is not specific to the environment or to toxins produced by *P. aeruginosa* but is instead the manifestation of a more generalized host response.

If the genetic mechanism underlying resistance consisted of a single stress response activated in both environments, the expected outcome would be for a strongly correlated and reciprocal resistant phenotype in the populations evolved in *P. aeruginosa* conditions. As shown here, evolution in the toxin environment did not confer any reciprocal increase in resistance to the infectious environment (Figure 2B). On the other hand, populations evolved under infection do show an increase in resistance to the toxin when compared to the ancestral population albeit to a lesser degree than the population directly selected for resistance to the toxin (Supplemental Figure 1).

At first glance this appears to suggest that evolution in infectious conditions does offer some protection against the toxin. However, both control populations, when evolved in a similar manner but with the OP50 strain of *E. coli* replacing *P. aeruginosa*, evolved the same level of resistance to the toxin as the long-term infection populations (Supplemental Figure 1c). This suggests that the increase in toxin resistance displayed by the infection evolved population is the result of adaptation to an effect that is not dependent on the presence of *P. aeruginosa*. Because all the evolved lines (but not the ancestor) showed increased toxin resistance, it seems likely that resistance to the toxin is a byproduct of general laboratory adaptation, potentially via chronic exposure to *E. coli*, which is a component of the propagation of every line except the population exposed to the alternating *P. aeruginosa* environments. This is consistent with evidence that the OP50 strain of *E. coli* is mildly pathogenic to *C. elegans* (So, et al., 2011). Like *P. aeruginosa*, OP50 is a gram-negative bacterium and thereby they have many genetic features in common, some of which have been shown to attenuate toxin-mediated virulence in *P. aeruginosa* mutants (Mahajan-Miklos, et al., 1999). For example, a bacterial gene, *MdoH*, identified to be important for full toxin pathogenicity by *P. aeruginosa* is also present in OP50 (Mahajan-Miklos, et al., 1999). On the other hand, the *P. aeruginosa* toxin produced in response to high osmolarity in this study, phenazine-1-carboxylic-acid (PCA) (Cezairliyan, et al., 2013), is not produced by OP50. This suggests that killing by PCA is a multifactorial process and that some of the sub-lethal pathogenic properties of *P. aeruginosa* may be shared with OP50. Consistent with this, the genes thought to be responsible for the presumed primary toxic compounds produced by *P. aeruginosa* in the infectious environment - phenazines (and in particular the phenazine pyocyanin) - are generally not present in the genus *Escherichia* and therefore are not found in OP50 (Pierson & Pierson, 2010; Williams, et al., 2010). Importantly, this generalized resistance to the toxin is not dependent on some important features of environment, such as osmolarity per se, although an exhaustive exploration of underlying causation here warrants further exploration. Unlike the resistance to toxin, the evolution of resistance to chronic infection was a much more specific response. Independent action of virulence factors is the best way to account for the lack of any correlated increase in infection resistance from other environments. Taken together these data suggest the genetic pathways underlying resistance to infection and toxin-mediated killing by *P. aeruginosa* are largely independent of one another with an overlap in resistance to the toxin-producing environment mediated by adaptation to the laboratory environment, most likely via OP50.

An additional perspective on the evolutionary independence of the genetic pathways for infection resistance and toxin resistance comes from examination of the population evolved under alternating infectious and toxin conditions. The alternating populations display no significant increase in resistance to infection but, despite having no exposure to OP50 during the period of experimental evolution, do have an increased resistance to the toxin environment relative to the ancestral population (Figure 2C). However, the level of resistance is significantly lower than the degree displayed by the populations evolved under infectious or control conditions (Supplemental Figure 1). Lack of evolution of resistance to infection is suggestive of an evolutionary antagonistic interaction between the infection and toxin resistance pathways. Additionally, evidence from assays in *C. elegans* demonstrates that the transcription of antibacterial immune effectors is selectively repressed during infection with the fungal pathogen *Candida albicans* and that production of an antimicrobial peptide required for defense against fungal pathogens during infection by a bacterial pathogen increases susceptibility (Pukkila-Worley, Mylonakis, & Ausubel, 2011; Engelmann, et al., 2011; Marsh, et al., 2011). This suggests that the worms are capable of distinguishing between pathogen damage types and coordinating responses among independent and antagonistic genetic pathways, but this interpretation is complicated by differences in the intensity of selection across the alternating environments (Pukkila-Worley, 2016). Because mortality was at 70% within 24 hours in the toxin-producing environment, non-resistant individuals exposed as L4s were likely to die before producing offspring whereas mortality in the infection environment reaches 50% around day 5, giving individuals exposed as L4 larvae ample time to produce offspring. The greater response to resistance to toxin could therefore in part be generated by greater intensity of selection in this environment, but only an antagonistic relationship can explain the lack of response to resistance to infection. Independent and asymmetric responses to selection for resistance to pathogens is consistent with asymmetric responses to selection for resistance to heat and oxidation stress within these same populations (Sikkink, et al., 2015).

This study suggests that *C. remanei* responds to virulence mechanisms of *P. aeruginosa* by at least two distinct genetic pathways. These two independent pathways are likely initiated in response to environment-specific factors through as-yet unknown interfaces. These pathways appear to interact in a somewhat antagonistic fashion. In the context of host-pathogen interactions and bacteriovore feeding systems, this may reflect strategies evolved by the host to balance energy investment in antimicrobial defenses while addressing the capabilities of pathogenic bacteria to evade detection and predation. It does, however, suggest that there may be some constraint on the evolution of generalized resistance phenotypes, at least in the short term.

## MATERIALS AND METHODS

### Bacterial Strains, Growth Conditions, and Media

The *P. aeruginosa* strain PA14 (Rahme, et al., 1995) and the *E. coli* strain OP50 were used. Strains were grown at 37°C in King’s broth (KB) (King, et al., 1954) or Luria-Bertani broth (LB) (Miller, 1972), respectively. PA14 was cultured on nematode growth medium lite (NGM lite) (US Biological catalog #N1005) for infection assays or peptone-glucose-sorbitol (PGS) media (1% Bacto-Peptone, 1% NaCl, 1% glucose, 1.7% Bacto-Agar, 0.15 M sorbitol) for toxin assays (Mahajan-Miklos, et al., 1999; Tan, et al., 1999). OP50 cultured on NGM lite was used for a low-osmolarity control and OP50 cultured on PGS was used for a high-osmolarity control.

### Nematode Strains

All *C. remanei* strains were maintained by standard culturing conditions on NGM lite agar with *E. coli* OP50 or *P. aeruginosa* PA14 as a food source at 20°C (Stiernagle, 1999). The strain used in this work was PX443, a derivative of 39 natural isolates collected from wood lice found in the same tract of forest detritus in the Koffler Scientific Reserve at Jokers Hill, King Township, Ontario. The isolates were intercrossed in a scheme designed to equalize the genomic contribution from each strain to generate PX443 (as described in Sikkink, et al., 2015).

### Bacterial Virulence Survival Assays

Both the toxin and infection assays were based on protocols described in Mahajan-Miklos et al. (1999). The toxin assay was conducted by spreading 60 μL of liquid from a PA14 culture grown for 18 hours in KB on 11 cm plates containing peptone-glucose-sorbitol medium. After the culture was spread, plates were incubated at 37°C for 24 hours then placed at room temperature for 2-4 hours. Approximately 200 age-synchronized L4 individuals per plate were placed on toxin-producing media for survival as well as PGS and NGM lite seeded with OP50 as controls and incubated at room temperature for 4 hours. The plates were then washed with S-basal buffer solution (Stiernagle, 1999), worms transferred onto NGM lite-OP50 plates, and surviving individuals counted 24 hours later. A total of three replicate plates per test condition were used.

The infection assay was conducted by spreading 20μL of liquid from a PA14 culture grown for 18 hours in KB on 3 cm plates containing NGM lite. The plates were then incubated at 37°C for 24 hours and placed at room temperature for 2-4 hours. A total of 90 age-synchronized L4 females per line were picked to said plates at a density of 3 females per plate. Resistance was scored as the fraction surviving after 5 days of exposure to the infectious PA14. An individual was considered dead when a nose-touch failed to elicit any movement.

### Experimental Evolution

#### Selection under toxin-producing conditions

Two population replicates of *C. remanei* (PX443) were grown on NGM lite plates with OP50 as a food source for 40 generations. This population was exposed to PA14 on PGS solid media for 4 hours every other generation upon reaching the L4 developmental stage. High-osmolarity controls were passaged in the same manner but with OP50 substituted for PA14. A subset of each population was frozen every fifth generation for later analysis.

#### Selection under chronic infection

Two population replicates of *C. remanei* (PX443) were grown on NGM lite plates alternating PA14 and OP50 as the food source every other generation for 40 generations. Low-osmolarity (lab-adapted) controls were passaged in the same manner but with OP50 substituted for PA14. A subset of each population was frozen every fifth generation for later analysis.

#### Selection in alternating environments

Two population replicates of *C. remanei* (PX443) were grown on NGM lite plates with PA14 as the only food source for 40 generations. This population was also exposed to PA14 on PGS solid media for 4 hours every other generation upon reaching the L4 stage of development. A subset of the population was frozen every fifth generation for later analysis.

### Statistical Analysis

Infection survival was calculated as the proportion of individuals surviving after five days of exposure. Individuals who had left the plates were removed from the analysis. Survival in the presence of toxin was calculated as the proportion of individuals surviving after 24 hours following removal from the toxin-producing environment. Survivorship was normalized to the number of individuals 24 hours after transfer from control (OP50) plates. Specific hypotheses were tested via categorical data analysis of weighted survival frequencies using JMP Pro10. Differences among groups was tested using the chi-square resulting from the appropriate likelihood-ratio test.

## Acknowledgements

The authors thank Karen Guillemin, John Willis, and Rose Reynolds for their supportive comments and helpful suggestions, and undergraduates Angela Uys and John Beckford for their contributions to this work.

## REFERENCES

Albertson RC, Cresko W, Detrich HW, Postlethwait JH (2009) Evolutionary mutant models for human disease. Trends Genet 25(2): 74–81.

Andersen EC, Gerke JP, Shapiro JA, Crissman JR, Ghosh R, Bloom JS, Félix M-A, Kruglyak L (2012) Chromosome-scale selective sweeps shape *Caenorhabditis elegans* genomic diversity. Nature Genetics 44(3): 285–290.

Arvanitis M, Glavis-Bloom J, Mylonakis E (2013) *C. elegans* for anti-infective discovery. Current Opinion in Pharmacology 13(5): 769–774.

Barriére A, Félix M-A (2005) Natural variation and population genetics of *Caenorhabditis elegans*. The *C. elegans* research community, WormBook 1551–8507 doi/10.1895/wormbook.1.43.1, http://www.wormbook.org

Boulin T, Hobert O (2012) From genes to function: The *C elegans* genetic toolbox. Wiley Interdiscip Rev Dev Biol 1(1): 114–137.

*C elegans* Deletion Mutant Consortium (2012) Large-scale screening for targeted knockouts in the *Caenorhabditis elegans* genome. G3 2(11): 1415–1425.

Cezairliyan B, Vinayavekhin N, Grenfell-Lee D, Yuen GJ, Saghatelian A, Ausubel FM (2013) Identification of *Pseudomonas aeruginosa* phenazines that kill *Caenorhabditis elegans*. PLoS Pathog 9(1): e1003101.

Cutter AD, Baird SE, Charlesworth D (2006) High nucleotide polymorphism and rapid decay of linkage disequilibrium in wild populations of *Caenorhabditis remanei*. Genetics 174(2): 901–913.

Engelmann I, Griffon A, Tichit L, Montañana-Sanchis F, Wang G, Reinke V, et al. (2011) A comprehensive analysis of gene expression changes provoked by bacterial and fungal infection in *C. elegans*. PLoS ONE 6(5): e19055.

Feinbaum RL, Urbach JM, Liberati NT, Djonovic S, Adonizio A, Carvunis A-R, et al. (2012) Genome-wide identification of *Pseudomonas aeruginosa* virulence-related genes using a *Caenorhabditis elegans* infection model. PLoS Pathog 8(7): e1002813. https://doi.org/10.1371/journal.ppat.1002813

Fierst JL, Willis JH, Thomas CG, Wang W, Reynolds RM, et al. (2015) Reproductive mode and the evolution of genome size and structure in *Caenorhabditis* nematodes. PLoS Genet 11(6): e1005323.

Gao AW, Sterken MG, uit de Bos J, van Creij J, Kamble R, et al. (2018) Natural genetic variation in *C. elegans* identified genomic loci controlling metabolite levels. Genome Research 28(9): 1296–1308.

Graustein A, Gaspar JM, Walters JR, Palopoli MF (2002) Levels of DNA polymorphism vary with mating system in the nematode genus *Caenorhabditis*. Genetics 161(1): 99–107.

Hahnel SR, Zdraljevic S, Rodriguez BC, Zhao Y, McGrath PT, Andersen EC (2018) Extreme allelic heterogeneity at a *Caenorhabditis elegans* beta-tubulin locus explains natural resistance to benzimidazoles. PLoS Pathog 14(10): e1007226.

Irazoqui JE, Troemel ER, Feinbaum RL, Luhachack LG, Cezairliyan BO, Ausubel FM (2010) Distinct pathogenesis and host responses during infection of *C. elegans* by *P. aeruginosa* and *S. aureus*. PLoS Pathog 6(7): e1000982. https://doi.org/10.1371/journal.ppat.1000982

Jovelin R, Ajie BC, Phillips PC (2003) Molecular evolution and quantitative variation for chemosensory behaviour in the nematode genus *Caenorhabditis*. Molecular Ecology 12(5): 1325–1337.

Kammenga JE, Phillips PC, De Bono M, Doroszuk A (2008) Beyond induced mutants: using worms to study natural variation in genetic pathways. Trends in Genetics 24(4): 178–185.

King EO, Ward MK, Raney DE (1954) Two simple media for the demonstration of pyocyanin and fluorescein. J Lab Clin Med 44: 301.

Kirienko NV, Cezairliyan BO, Ausubel FM, Powell JR (2014) *Pseudomonas aeruginosa* PA14 pathogenesis in *Caenorhabditis elegans*. In Pseudomonas: Methods and Protocols (Vol. 1149, Methods in Molecular Biology, pp. 653–669). TOTOWA, NJ: HUMANA PRESS. doi: 10.1007/978-1-4939-0473-0 50.

Mahajan-Miklos S, Tan MW, Rahme LG, Ausubel FM (1999) Molecular mechanisms of bacterial virulence elucidated using a *Pseudomonas aeruginosa Caenorhabditis elegans* pathogenesis model. Cell 96(1): 47–56.

Marsh EK, van den Berg MCW, May RC (2011) A two-gene balance regulates *Salmonella typhimurium* tolerance in the nematode *Caenorhabditis elegans*. PLoS ONE 6(3): e16839.

Miller JH (1972) Experiments in molecular genetics. Cold Spring Harbor Laboratory, Cold Spring Harbor, New York.

Morran LT, Schmidt OG, Gelarden IA, Parrish RC, Lively CM (2011) Running with the Red Queen: Host-parasite coevolution selects for biparental sex. Science 333(6039): 216–218.

Noble LM, Chelo I, Guzella T, Afonso B, Riccardi DD, et al. (2017) Polygenicity and epistasis underlie fitness-proximal traits in the *Caenorhabditis elegans* multiparental experimental evolution (CeMEE) panel. Genetics 207(4): 1663–1685.

O’Callaghan D, Vergunst A (2010) Non-mammalian animal models to study infectious disease: worms or fly fishing? Current Opinoin in Microbiology 13(1): 79–85.

Phillips PC (2012) Self-fertilization sweeps up variation in the worm genome. Nature Genetics 44(3): 237–238.

Pierson III LS, Pierson EA (2010) Metabolism and function of phenazines in bacteria: impacts on the behavior of bacteria in the environment and biotechnological processes. Appl Microbiol Biotechnol 86(6): 1659–1670.

Pukkila-Worley R, Ausubel FM, Mylonakis E (2011) *Candida albicans* infection of *Caenorhabditis elegans* induces antifungal immune defenses. PLoS Pathog 7(6): e1002074

Pukkila-Worley R (2016) Surveillance immunity: an emerging paradigm of innate defense activation in *Caenorhabditis elegans*. PLoS Pathog 12(9): e1005795.

Rahme LG, Stevens EJ, Wolfort SF, Shao J, Tompkins RG, Ausubel FM (1995) Common virulence factors for bacterial pathogenicity in plants and animals. Science 268(5219): 1899–1902.

Rockman MV (2012) The QTN program and the alleles that matter for evolution: all that’s gold does not glitter. Evolution 66(1): 1–17.

SchulenburgH, EwbankJJ (2004) Diversity and specificity in the interaction between Caenorhabditis elegans and the pathogen Serratia marcescens. BMC Evolutionary Biology 4:49.

Schulenburg H, Boehnisch C (2008) Diversification and adaptive sequence evolution of *Caenorhabditis* lysozymes (Nematoda: Rhabditidae). BMC Evolutionary Biology 8:114.

Sikkink KL, Reynolds RM, Cresko WA, Phillips PC (2015) Environmentally induced changes in correlated responses to selection reveal variable pleiotropy across a complex genetic network. Evolution 69(5): 1128–1142.

Slowinski SP, Morran LT, Parrish RC, Cui ER, Bhattacharya A, Lively CM, Phillips PC (2016) Coevolutionary interactions with parasites constrain the spread of self-fertilization into outcrossing host populations. Evolution 70(11): 2632–2639.

So SH, Tokumaru T, Miyahara K, Ohshima Y (2011) Control of lifespan by food bacteria, nutrient limitation and pathogenicity of food in *C. elegans*. Mechanisms of Ageing and Development 132(4): 210–212.

Stiernagle T (1999) Maintenance of *C. elegans*. In C. elegans: a practical approach (Vol. 213, Practical Approach Series, pp. 51–67). NEW YORK, NY: OXFORD UNIVERSITY PRESS.

Tan MW, Mahajan-Miklos S, Ausubel FM (1999) Killing of *Caenorhabditis elegans* by *Pseudomonas aeruginosa* used to model mammalian bacterial pathogenesis. PNAS 96(2): 715–720.

Tan MW, Rahme LG, Sternberg JA, Tompkins RG, Ausubel FM (1999) *Pseudomonas aeruginosa* killing of *Caenorhabditis elegans* used to identify *P. aeruginosa* virulence factors. PNAS 96(5): 2408–2413.

Teotónio H, Estes S, Phillips PC, Baer CF (2017) Experimental evolution with *Caenorhabditis* nematodes. Genetics 206(2): 691–716.

Wang Y, Wilks JC, Danhorn T, Ramos I, Croal L, Newman DK (2011) Phenazine-1-Carboxylic Acid promotes bacterial biofilm development via ferrous iron acquisition. Journal of Bacteriology 193(14): 3606–3617.

Williams KP, Gillespie JJ, Sobral BWS, Nordberg EK, Snyder EE, Shallom JM, Dickerman AW (2010) Phylogeny of gammaproteobacteria. Journal of Bacteriology 192(9): 2305–2314.

