## Supplemental File for "Experimental evolution of independent genetic pathways for resistance to *Pseudomonas aeruginosa* pathogenicity within the nematode *Caenorhabditis remanei*"

* † ‡

†

Log Fold Change in Survival

| Toxin Environment | Toxin Adapted | Infection Adapted | Alternating Adaptation | High Osmolarity Control | Low Osmolarity Control |
| --- | --- | --- | --- | --- | --- |
| 1 | 0.483676 | 0.410305 | 0.268773 | 0.397751 | 0.377328 |

Supplemental Figure 1. Log fold survival relative to ancestor of all evolved populations tested in the toxin-producing environment ($X_{4}^{2}$ = 241.144, p < 0.0001; * = $X_{3}^{2}$ = 84.639, p < 0.0001; † = $X_{1}^{2}$ = 67.059, p < 0.0001; ‡ = $X_{2}^{2}$ = 3.785, p = 0.15).

Log Fold Change in Survival

| Infection Environment | Toxin Adapted | Infection Adapted | Alternating Adaptation | High Osmolarity Control | Low Osmolarity Control |
| --- | --- | --- | --- | --- | --- |
| 1 | 0.0429839 | 0.166246 | -0.0599517 | -0.14432 | -0.152619 |

Supplemental Figure 2. Log fold survival relative to ancestor of all evolved populations tested in the infection environment ($X_{4}^{2}$ = 59.501, p < 0.0001).

Supplemental Table 1. Pairwise chi-square analysis of all replicates. Shaded cells = toxin assay, no shading = infection assay.

| Toxin Evolved 1 | Toxin Evolved 2 | Infection Evolved 1 | Infection Evolved 2 | Alternating Evolved 1 | Alternating Evolved 2 | High Osmolarity Control 1 | High Osmolarity Control 2 | Low Osmolarity Control 1 | Low Osmolarity Control 2 | Ancestor |  |
| --- | --- | --- | --- | --- | --- | --- | --- | --- | --- | --- | --- |
|  | Χ^2^_1_ = 0.2, p = 0.6335 | Χ^2^_1_ = 6.5, p = 0.0107 | Χ^2^_1_ = 1.6, p = 0.2085 | Χ^2^_1_ = 0.7, p = 0.4031 | Χ^2^_1_ = 5.1, p = 0.0245 | Χ^2^_1_ = 8.4, p = 0.0037 | Χ^2^_1_ = 5.3, p = 0.0214 | Χ^2^_1_ = 5.8, p = 0.0162 | Χ^2^_1_ = 7.1, p = 0.0076 | Χ^2^_1_ = 1.2, p = 0.2838 | Toxin Evolved 1 |
| Χ^2^_1_ = 1.2, p = 0.2743 |  | Χ^2^_1_ = 9.9, p = 0.0017 | Χ^2^_1_ = 3.2, p = 0.0718 | Χ^2^_1_ = 0.1, p = 0.7140 | Χ^2^_1_ = 3.4, p = 0.0655 | Χ^2^_1_ = 6.4, p = 0.0112 | Χ^2^_1_ = 3.7, p = 0.0546 | Χ^2^_1_ = 4.1, p = 0.0438 | Χ^2^_1_ = 5.2, p = 0.0226 | Χ^2^_1_ = 0.4, p = 0.5404 | Toxin Evolved 2 |
| Χ^2^_1_ = 37.3, p < 0.0001 | Χ^2^_1_ = 52.5, p < 0.0001 |  | Χ^2^_1_ = 1.8, p = 0.1820 | Χ^2^_1_ = 12.7, p = 0.0004 | Χ^2^_1_ = 25.5, p < 0.0001 | Χ^2^_1_ = 30.9, p < 0.0001 | Χ^2^_1_ = 23.8, p < 0.0001 | Χ^2^_1_ = 26.0, p < 0.0001 | Χ^2^_1_ = 30.1, p < 0.0001 | Χ^2^_1_ = 14.5, p = 0.0001 | Infection Evolved 1 |
| Χ^2^_1_ = 14.2, p = 0.0002 | Χ^2^_1_ = 23.0, p < 0.0001 | Χ^2^_1_ = 2.9, p = 0.0916 |  | Χ^2^_1_ = 4.8, p = 0.0281 | Χ^2^_1_ = 13.5, p = 0.0002 | Χ^2^_1_ = 18.1, p < 0.0001 | Χ^2^_1_ = 13.0, p = 0.0003 | Χ^2^_1_ = 14.2, p = 0.0002 | Χ^2^_1_ = 16.9, p < 0.0001 | Χ^2^_1_ = 6.0, p = 0.0146 | Infection Evolved 2 |
| Χ^2^_1_ = 99.6, p < 0.0001 | Χ^2^_1_ = 119.1, p < 0.0001 | Χ^2^_1_ = 30.7, p < 0.0001 | Χ^2^_1_ = 38.7, p < 0.0001 |  | Χ^2^_1_ = 2.3, p = 0.1326 | Χ^2^_1_ = 5.0, p = 0.0256 | Χ^2^_1_ = 2.6, p = 0.1070 | Χ^2^_1_ = 2.9, p = 0.0902 | Χ^2^_1_ = 3.8, p = 0.0510 | Χ^2^_1_ = 0.1, p = 0.8023 | Alternating Evolved 1 |
| Χ^2^_1_ = 115.5, p < 0.0001 | Χ^2^_1_ = 138.2, p < 0.0001 | Χ^2^_1_ = 35.5, p < 0.0001 | Χ^2^_1_ = 42.8, p < 0.0001 | Χ^2^_1_ = 0.2, p = 0.6989 |  | Χ^2^_1_ = 0.7, p = 0.4063 | Χ^2^_1_ = 0.1, p = 0.8056 | Χ^2^_1_ = 0.1, p = 0.7955 | Χ^2^_1_ = 0.2, p = 0.6517 | Χ^2^_1_ = 1.6, p = 0.2098 | Alternating Evolved 2 |
| Χ^2^_1_ = 22.8, p < 0.0001 | Χ^2^_1_ = 34.2, p < 0.0001 | Χ^2^_1_ = 0.9, p = 0.3505 | Χ^2^_1_ = 0.6, p = 0.4481 | Χ^2^_1_ = 34.3, p < 0.0001 | Χ^2^_1_ = 38.4, p < 0.0001 |  | Χ^2^_1_ = 0.3, p = 0.5972 | Χ^2^_1_ = 0.3, p = 0.5801 | Χ^2^_1_ = 0.2, p = 0.6821 | Χ^2^_1_ = 4.0, p = 0.0457 | High Osmolarity Control 1 |
| Χ^2^_1_ = 39.3, p < 0.0001 | Χ^2^_1_ = 54.0, p < 0.0001 | Χ^2^_1_ = 0.4, p = 0.5356 | Χ^2^_1_ = 4.4, p = 0.0353 | Χ^2^_1_ = 22.5, p < 0.0001 | Χ^2^_1_ = 24.7, p < 0.0001 | Χ^2^_1_ = 2.0, p = 0.1541 |  | Χ^2^_1_ = 0.0, p = 1.0000 | Χ^2^_1_ = 0.03, p = 0.8709 | Χ^2^_1_ = 2.0, p = 0.1663 | High Osmolarity Control 2 |
| Χ^2^_1_ = 43.9, p < 0.0001 | Χ^2^_1_ = 58.5, p < 0.0001 | Χ^2^_1_ = 2.1, p = 0.1508 | Χ^2^_1_ = 7.4, p = 0.0066 | Χ^2^_1_ = 14.0, p = 0.0002 | Χ^2^_1_ = 14.4, p = 0.0002 | Χ^2^_1_ = 4.5, p = 0.0344 | Χ^2^_1_ = 0.7, p = 0.4137 |  | Χ^2^_1_ = 0.03, p = 0.8641 | Χ^2^_1_ = 2.1, p = 0.1455 | Low Osmolarity Control 1 |
| Χ^2^_1_ = 27.3, p < 0.0001 | Χ^2^_1_ = 38.4, p < 0.0001 | Χ^2^_1_ = 0.1, p = 0.8018 | Χ^2^_1_ = 2.6, p = 0.1082 | Χ^2^_1_ = 19.3, p < 0.0001 | Χ^2^_1_ = 19.9, p < 0.0001 | Χ^2^_1_ = 1.0, p = 0.3284 | Χ^2^_1_ = 0.1, p = 0.8040 | Χ^2^_1_ = 0.9, p = 0.3470 |  | Χ^2^_1_ = 2.9, p = 0.0887 | Low Osmolarity Control 2 |
| Χ^2^_1_ = 265.4, p < 0.0001 | Χ^2^_1_ = 295.1, p < 0.0001 | Χ^2^_1_ = 159.2, p < 0.0001 | Χ^2^_1_ = 152.0, p < 0.0001 | Χ^2^_1_ = 33.4, p < 0.0001 | Χ^2^_1_ = 46.6, p < 0.0001 | Χ^2^_1_ = 152.1, p < 0.0001 | Χ^2^_1_ = 129.6, p < 0.0001 | Χ^2^_1_ = 98.9, p < 0.0001 | Χ^2^_1_ = 103.4, p < 0.0001 |  | Ancestor |
